# A Laboratory-Adapted and a Clinical Isolate of Dengue Virus Serotype 4 Differently Impact Aedes aegypti Life-History Traits Relevant to Vectorial Capacity

**DOI:** 10.1101/2024.06.07.597920

**Authors:** Mariana Maraschin, Diego Novak, Valdorion José Klein Junior, Lucilene W Granella, Luiza J. Hubner, Athina R Medeiros, Tiago Graf, Guilherme Toledo-Silva, Daniel S Mansur, José Henrique M Oliveira

## Abstract

1

Dengue virus cases are on the rise globally, and strategies to reduce new infections by controlling its primary vector, the mosquito *Aedes aegypti*, represent a promising biotechnological approach. However, the interaction between virus serotypes and genotypes with *Aedes aegypti* is poorly characterized at the molecular level, as well as in terms of life-history traits related to vector capacity and mosquito disease tolerance. Here, we infected *Aedes aegypti* with two philogenetically distant strains of Dengue virus serotype 4 genotype II: a laboratory-adapted strain, DENV4 - TVP/360, and a recent clinical strain isolated from southern Brazil, DENV4 - LRV 13/422. These strains, which exhibit 26 amino acid differences in their sequences, have been shown in to produce distinct immune phenotypes in vertebrate cells. Here, we assessed various life-history traits of *Aedes aegypti*, including mortality, fecundity, fertility, and induced flight capacity, as well as vector competence-related parameters such as infection intensity and prevalence, following exposure to different viral concentrations. We found that each viral strain differentially affected mosquito traits. While neither strain significantly reduced mosquito lifespan, *Aedes aegypti* infection prevalence was influenced by the initial dose of DENV4 - TVP/360. This laboratory-adapted strain also enhanced mosquito induced-flight capacity at early (24 hours post-infection - DPI) and late (21 DPI) time points. On the other hand, the recent clinical isolate, DENV4 - LRV 13/422, specifically reduced *Aedes aegypti* fertility. A better understanding of how different arbovirus strains influence mosquito life-history traits connected to disease spread will be critical in public health efforts to mitigate arbovirus outbreaks that are focused on the mosquito vector.

**Highlights:**

- The DENV4 strains TVP/360 and LRV 13/422 have 26 amino acid differences in their sequences.
- *Aedes aegypti* infection prevalence is influenced by the initial dose of DENV4 - TVP/360 strain but not DENV4 - LRV 13/422.
- Fertility is specifically reduced by DENV4 - LRV 13/422.
- Induced-flight activity is enhanced at 1 and 21 days post-infection with DENV4 - TVP/360 strain.

## 3 Introduction

The ability of hosts to sustain infection without significant impact on their health and fitness is called disease tolerance, a key feature of vector competence and pathogen transmission by mosquitoes (Oliveira et al. 2020). *Aedes aegypti* is predominantly tolerant to the arboviruses they transmit, such as Dengue and Zika (Lambrechts and Saleh 2019; Magistrado, El-Dougdoug, and Short 2023). Several life-history traits of mosquitoes, such as survival, reproductive output, and dispersal influence arbovirus transmission (Cator et al. 2020) and can be used to quantify mosquito disease tolerance (Maraschin et al. 2023). Well-defined mosquito health metrics are critical for a better understanding of the molecular mechanisms that enable *Aedes aegypt*i to sustain a chronic arbovirus infection during its life cycle. Identifying and characterizing such mechanisms and how their manipulation changes transmission is fundamental to developing strategies to block vector competence and arbovirus epidemics (Chandrasegaran et al. 2020; Wu et al. 2020).

In this manuscript, we evaluated the impact of two Dengue virus serotype 4 genotype II strains (DENV4) in different life-history traits of *Aedes aegypti*. The strains are a laboratory-adapted reference strain DENV4 - TVP/360 (GenBank - KU513442) and a clinical isolate from a patient presenting high serum viral load in southern Brazil, DENV4 - LRV 13/422 (GenBank - KU513441) (Sarathy et al. 2015; Kuczera et al. 2016). They were chosen because they present different *in vitro* infectivity rates in vertebrate and mosquito cells and immunomodulatory capacities in vertebrate cells. We hypothesized that the two strains would differentially impact *Aedes aegypti* life-history traits under laboratory conditions. We focused on parameters that are directly linked to mosquito health and fitness, such as survival, fecundity, and induced-flight activity (INFLATE), a proxy for dispersion (Gaviraghi and Oliveira 2020). We found that the prevalence of infection differed between strains, with only the laboratory-adapted (DENV4 - TVP/360) being highly dependent on the initial virus titer. The infection did not significantly alter the day of mosquito death or fecundity (eggs/female) at the population level but reduced *Aedes aegypti* fertility (% hatching) specifically in the clinical isolate DENV4 - LRV 13/422. Interestingly, the laboratory-adapted DENV4 TVP/360 strain significantly and specifically enhanced the induced-flight activity (INFLATE) of *Aedes aegypti* at early (1-day post-infection - DPI) and late (21 DPI) time points. To modulate vector disease tolerance and reduce disease transmission, we will need to improve our ability to quantify the mosquito health status during infection (Shirasu-Hiza and Schneider 2007; Torres et al. 2016; Louie et al. 2016) and understand the molecular basis of strain-specific effects in critical life-history traits, such as survival and reproductive output as well as underexplored mosquito phenotypes connected to vectorial capacity, such as the induced-flight activity.

## 4 Material and Methods

### 4.1 Mosquitoes: *Aedes aegypti*

(Red Eye strain) were reared and maintained under standard conditions as described previously (Jose Henrique M. Oliveira et al. 2011; José Henrique M. Oliveira et al. 2017) at the Federal University of Santa Catarina, Brazil, with a 12-hour light/dark cycle at 28°C and 70–80% relative humidity. Larvae were maintained in filtered and dechlorinated tap water and fed with powdered dog chow (2-3 additions during the larval stage without changing the water). Adults were provided a 10% sucrose solution *ad libitum* while housed in 7.3-liter plastic cages (21 cm diameter x 25 cm height) with ∼ 200 - 300 mosquitoes per cage. Infections were performed in females aged between 3 and 7 days post-adult eclosion, which were used in all assays and maintained under the same light-dark cycle at 28°C and 80% relative humidity. Additionally, for egg production to maintain the insectarium, females were fed artificially using a water-jacketed glass artificial feeder and a parafilm membrane containing peripheral human blood (collected in the presence of EDTA) and 1mM ATP as a phagostimulant. Informed consent was obtained from all blood donors. This protocol was approved by the Federal University of Santa Catarina (UFSC) - CAAE: 89894417.8.0000.0121.

### 4.2 Dengue virus serotype 4 stock preparations

We used Dengue virus 4 strain strain TVP/360 – GenBank: KU513442, hereafter called DENV4/TVP, and Dengue virus 4 strain LRV13/422 - GenBank: KU513441, hereafter called DENV4/LRV (Kuczera et al. 2016). Virus strains were kindly provided by Dr Claudia Nunes Duarte dos Santos from Instituto Carlos Cagas - Fiocruz Paraná, Brazil. *Aedes albopictus* mosquito cells of the C6/36 lineage were maintained and propagated in 1X L-15 medium with a pH of 7.6, supplemented with 5% SFB, 1% P/S, and 0.26% tryptone, at a temperature of 28°C in a Biochemical Oxygen Demand (B.O.D.) oven. The cells were seeded at 80% confluency (3 x 10^7 cells) in a 150 cm^2 bottle, which was kept at 28°C overnight. The following day, the cells were infected with a Multiplicity of Infection (MOI) of 0.01 for 5 days to produce DENV4/TVP or DENV4/LRV. The cell supernatant, containing viral particles, was collected, centrifuged at 460 x g for 10 minutes at 4°C, aliquoted, and stored at −80°C. The plaque assay was performed to determine the viral titer.

### 4.3 Phylogenetic analysis

A comparative phylogenetic analysis between LRV 13/422 and TVP/360 strains was performed by using augur/auspice as available in Nextstrain (Hadfield et al. 2018). Initially, all DENV4 genomes with more than 70% coverage compared to the reference strain (NC_002640) were retrieved from NCBI Virus. Sequences were then aligned with MAFFT (Katoh and Standley 2013) and visually inspected in Aliview (Larsson 2014). A maximum likelihood phylogenetic tree was constructed with IQ-TREE (Nguyen et al. 2015) and the best substitution model was inferred by the model testing function. Branch support was calculated with SH-aLRT in 1000 pseudoreplicates. The phylogenetic tree was then visualized in FigTree (https://github.com/rambaut/figtree), revealing that both LRV 13/422 and TVP360 strains and the DENV4 reference genome belonged to genotype 2. We then reconstructed the ancestral sequences of this genotype and translated the mutations using augur. The sequence AF326573 was used as the root sequence since NC_002640 was derived from an engineered strain (AF326825) and AF326573 was the natural isolation from which AF326825 originated (Durbin et al. 2001). Auspice was used to visualize and stylize the tree and to extract LRV 13/422 and TVP/360 mutation path from the root sequence.

### 4.4 *Aedes aegypti* infection with Dengue virus serotype 4

For mosquito infection experiments, human blood was collected from healthy donors (UFSC - CAAE: 89894417.8.0000.0121) using tubes containing EDTA reagent. To isolate red blood cells (RBCs), 1 mL of blood was centrifuged at 6,400 rpm for 4 minutes at room temperature. Following centrifugation, the serum was discarded, and the cells were gently washed twice with sterile 1X PBS solution (GIBCO). *Aedes aegypti* females were maintained with access to water but were fasted on sucrose for 18-24 hours before being offered a meal containing a 1:1 mixture of RBC cells and L-15 medium containing either DENV4 TVP/360 virus or DENV4 LRV13/422. The control group (Mock) received a 1:1 mixture of RBC and C6/36 cell supernatant. All groups included ATP, pH 7.4, at a final concentration of 1 mM as a phagostimulant. These solutions were presented to the females using artificial glass feeders heated by water at 37°C for approximately 1 hour. Subsequently, mosquitoes were anesthetized using cold (−20°C), and only fully engorged females were separated into small cages and kept in a B.O.D. incubator at 28°C until the experiment concluded. Each cage housed 20 mosquitoes and was supplied with a 10% sucrose solution soaked in cotton, offered *ad libitum*.

### 4.5 Plaque assays

Vero cells were used to quantify DENV4 TVP/360 and Vero E6 cells to quantify DENV4 LRV 13/422 based on plaque optimization for each viral strain. Cells were seeded at a density of 5 x 10^4 cells/well and 1 x 10^5 cells/well, respectively, in a 24-well plate in DMEM F-12 supplemented with 5% FBS, 1% penicillin/streptomycin, 1% glutamine (1X complete DMEM) maintained at 37°C and 5% CO2. The following day, the mosquito samples were thawed and underwent a decontamination process, which involved immersing each mosquito in 1 x 45” in 70% alcohol, followed by 1 x 45” in 1% hypochlorite, another round of 1 x 45” in 70% alcohol, and finally 1 x 45” in 0.9% sterile saline. Subsequently, each mosquito was transferred to a 1.5 mL tube containing 200 µL of complete DMEM F-12 medium (as defined above) and kept on ice. The mosquitoes were then macerated individually using a manual vortex with a sterile pestle dedicated to each sample. The homogenate was then centrifuged at 3,200 × g for 5 minutes at 4°C. Each mosquito homogenate was diluted (ranging from 10^-1 to 10^-5) in DMEM F-12 - 1X complete medium, added to Vero cells, and incubated in 200 µL of each dilution for 60 minutes at 37°C and 5% CO2. The same procedure was performed for the control group (Mock) using 200 µL of complete DMEM F-12 medium. After incubation, the medium containing the viral dilutions was removed, and 700 µL of semi-solid medium containing DMEM 2X supplemented with 1% fetal bovine serum, 1% P/S, and 1,6% carboxymethylcellulose was added. The plates were then incubated for 5 days at 37°C and 5% CO2. Following incubation, the cells were fixed in PFA 3% for 20 minutes, stained in 1% crystal violet, and counted.

### 4.6 Survival curves

All infections were performed in female mosquitoes between 4 and 6 days following adult emergence. Infected females were cold-anesthetized immediately after feeding and transferred to cardboard cups with a density of 20 fully engorged females per cup (maximum capacity of 470 mL - 10 cm height x 9 cm diameter). *Ad libitum* access to a 10% sucrose solution was provided through cotton pads placed on top of a woven mesh, which were replaced 2–3 times weekly. Survival rates were monitored daily, six times a week, until all insects within the cups died. The survival cages were maintained in the insectary at a temperature of 28°C (±10%) and humidity of 80% (±10%). Survival data presented represents pooled results from a minimum of 2 independent experiments.

### 4.7 *Aedes aegypti* fecundity and fertility

Female mosquitoes were artificially fed with whole blood, mock, or blood supplemented with DENV4-TPV/360 or DENV4-LRV 13/422 as detailed in item 4.4. Fully engorged females were cold-anesthetized and individually housed in cages containing a dark plastic cup with water and filter paper to allow egg deposition. Fecundity was scored 5 days following the meal by counting the number of eggs deposited per female. Collected eggs were allowed to rest in insectary conditions for 7 days when they were submerged in water containing filtered and dechlorinated tap water plus powdered dog chow to allow larval development. Fertility was assessed 3 days post eclosion by counting the percentage of ecloded larvae.

### 4.8 *Aedes aegypti* Induced-flight activity (INFLATE)

The protocol was adapted from (Gaviraghi and Oliveira 2020). A rectangular plastic cage measuring 19 cm x 20 cm was divided into four equal quadrants, in addition to the base. Each quadrant was assigned a value ranging from zero, the lowest (base), to four, the highest. Five mosquitoes were quickly anesthetized at a temperature of −20°C and introduced into the cage, where they remained for 10 minutes, acclimating to the experimental conditions in the insectary temperature and humidity. To begin the INFLATE test, a mechanical stimulus was applied to the cage, consisting of lifting it about 20 cm above the surface and gently tapping it, causing the mosquitoes to detach and fall to the base of the cage. The cage was kept stable for 1 minute, during which time the mosquitoes initiated flight activity. The number of mosquitoes that landed in each quadrant of the cage was visually recorded. Mosquitoes still flying after one minute were assigned the highest value (four). Mosquitoes landing on the line between quadrants received the value of the upper quadrant. Then, a 2 minute rest period was observed, and the process was repeated. Each repetition generated a value called the “inflate value,” calculated by multiplying the value of each quadrant by the number of mosquitoes landing in it. These values were then summed and divided by the total number of mosquitoes in the cage (five). This process was repeated 10 times, corresponding to a technical replicate (n=10). The average of the values from the 10 technical replicates is referred to as the “Inflate Index.” The protocol was conducted with infected and non-infected mosquitoes (control group, mock). Ultimately, a biological replicate was obtained (n=1). In Figure 4, each dot represents 1 biological replicate (consisting of 5 mosquitoes assayed 10 times).

### 4.9 Statistical analysis

Infection intensity, day of death, eggs per female, percentage of eclosion, and INFLATE index were tested for normality. For data that did not follow a normal distribution, non-parametric tests such as the Kruskal-Wallis test were performed, as indicated in the figure legends, for side-by-side comparisons or Dunn’s multiple comparisons test. Infection prevalence was analyzed using a contingency table (Yes - infected vs No - uninfected) and Fisher’s exact test. Survival curves were analyzed with a Log-rank (Mantel-Cox) test. Statistical analysis was conducted using GraphPad Prism.

## 5 Results

### Sequence differences between DENV4 - TVP/360 and DENV4 - LRV 13/422 sequences

We compared the sequences of the laboratory-adapted (DENV4 - TVP/360) and the clinical isolate (DENV4 - LRV 13/422) strains of DENV4 with a reference strain, DENV4/814669, obtained from the Dominican Republic in 1981 (Mackow et al. 1987; Durbin et al. 2001). Both strains were phylogenetically distant (Supplementary Figure 1) with the clinical isolate (LRV 13/422) exhibiting 24 unique amino acids (AA) substitutions compared to the reference sequence DENV4/814669, while the laboratory-adapted (TVP/360) strain exhibited 2 unique AA substitutions, compared to DENV4/814669 (Figure 1 and Table 1). Three mutations were shared by both strains, S2H, in the precursor Membrane glycoprotein (prM), K14Q, in the non-structural protein 3 (NS3), and R23K in the non-structural protein 5 (NS5). The 2 exclusive amino acid substitutions of DENV4 - TVP/360 strain were located in the non-structural protein NS4B, being phylogenetically closer to the reference strain, DENV4/814669, than the virus isolated from southern Brazil in 2013 (DENV4 - LRV 13/422). Most of the non-synonymous mutations of the Brazilian strain (LRV 13/422) were present in the envelope protein (E), with 6 AA substitutions; non-structural protein 1 (NS1) with 5 AA substitutions; non-structural protein 2A (NS2A), with 5 AA substitutions and the non-structural protein 5 (NS5), with 6 AA substitutions. Interestingly, these proteins are critical for flavivirus infection, replication, and dissemination into mosquito tissues (Erb et al. 2010; J. Liu et al. 2016; Y. Liu et al. 2017; Saraiva et al. 2018; Cheng et al. 2021; Besson et al. 2022).

**Figure 1.**
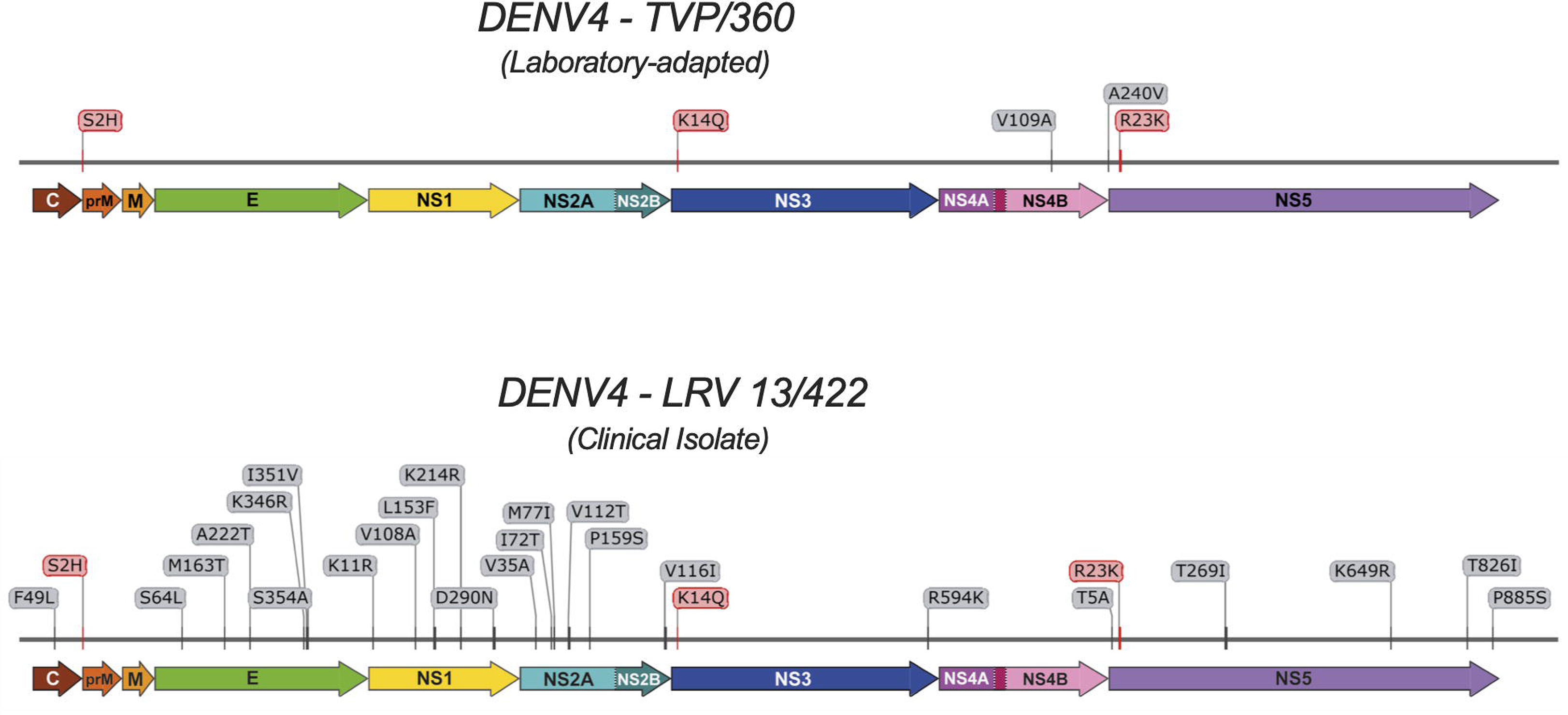
DENV4 Amino acid Sequence. Differences between strains compared to reference strain DENV4/814669 - GenBank: AF326573.1. Aminoacid (AA) substitutions highlighted in grey represent strain-specific mutations. Shared AA substitutions between DENV4 - TVP/360 and DENV4 - LRV 13/422 are highlighted in red.

**Table 1.** Aminoacid differences between DENV4 strains used. . Bold substitutions are present in both strains compared to the reference strain DENV4/814669 (GenBank: AF326573.1).

| DENV4 Genes | DENV4 - TVP/360 strain | DENV4 - LRV13/422 strain |
| --- | --- | --- |
| C |  | F49L |
| prM | **S2H** | **S2H** |
| E |  | S64L, M163T, A222T, K346R, I351V, S354A |
| NS1 |  | K11R, V108A, L153F, K214R, D290N |
| NS2A |  | V35A, I72T, M77I, V112T, P159S |
| NS2B |  | V116I |
| NS3 | **K14Q** | **K14Q**, R594K |
| NS4A |  |  |
| NS4B | V109A, A240V |  |
| NS5 | **R23K** | T5A, **R23K**, T269I, K649R, T826I, P885S |

### Infection dynamics of DENV4 strains differ in *Aedes aegypti*

To explore how the laboratory-adapted (DENV4 - TVP/360) and the clinical isolate (DENV4 - LRV 13/422) interact with *Aedes aegypti* over time, we infected mosquitoes with two doses of each strain and measured virus titers in single whole-body mosquitoes at 0, 7, 14 and 21 days following the infectious blood meal (Figure 2A and 2B). We used the maximum available dose for each strain and a 1/10 dilution. The infectious particle virus concentration (input in Figure 2B) is shown in red and expressed as plaque-forming units (PFU) per microliter of blood offered to mosquitoes (PFU/ul of blood). Both strains were able to infect and replicate in *Aedes aegypti* (Figure 2B). We were interested in two readouts, the infection prevalence, defined as the percentage of infected mosquitoes, and infection intensity, defined as the amount of infectious particles per mosquito. Figure 2C shows that despite a difference in input dose varying by a factor of 100X (from TVP - higher dose to LRV - lower dose), infection intensity had a minor impact on virus titer per mosquito. However, this range of input doses differentially influenced infection prevalence depending on the virus strain, with the laboratory-adapted strain (DENV4 - TVP/360) being highly influenced by virus input, as expected based on several mosquito-DENV studies (Nguyet et al. 2013; Duong et al. 2015; Novelo et al. 2019). However, the clinical isolate (DENV4 - LRV 13/422) exhibited a relatively low infection prevalence (∼40%), irrespective of the input dose (Figure 2C). This result suggests that factors determining infection prevalence, also known as the midgut infection barrier, might be more relevant to vector competence than immune resistance factors that decrease virus replication inside mosquito tissues once the infection is already established (Bennett et al. 2002; Souza-Neto, Powell, and Bonizzoni 2019; Hodoameda et al. 2024).

**Figure 2.**
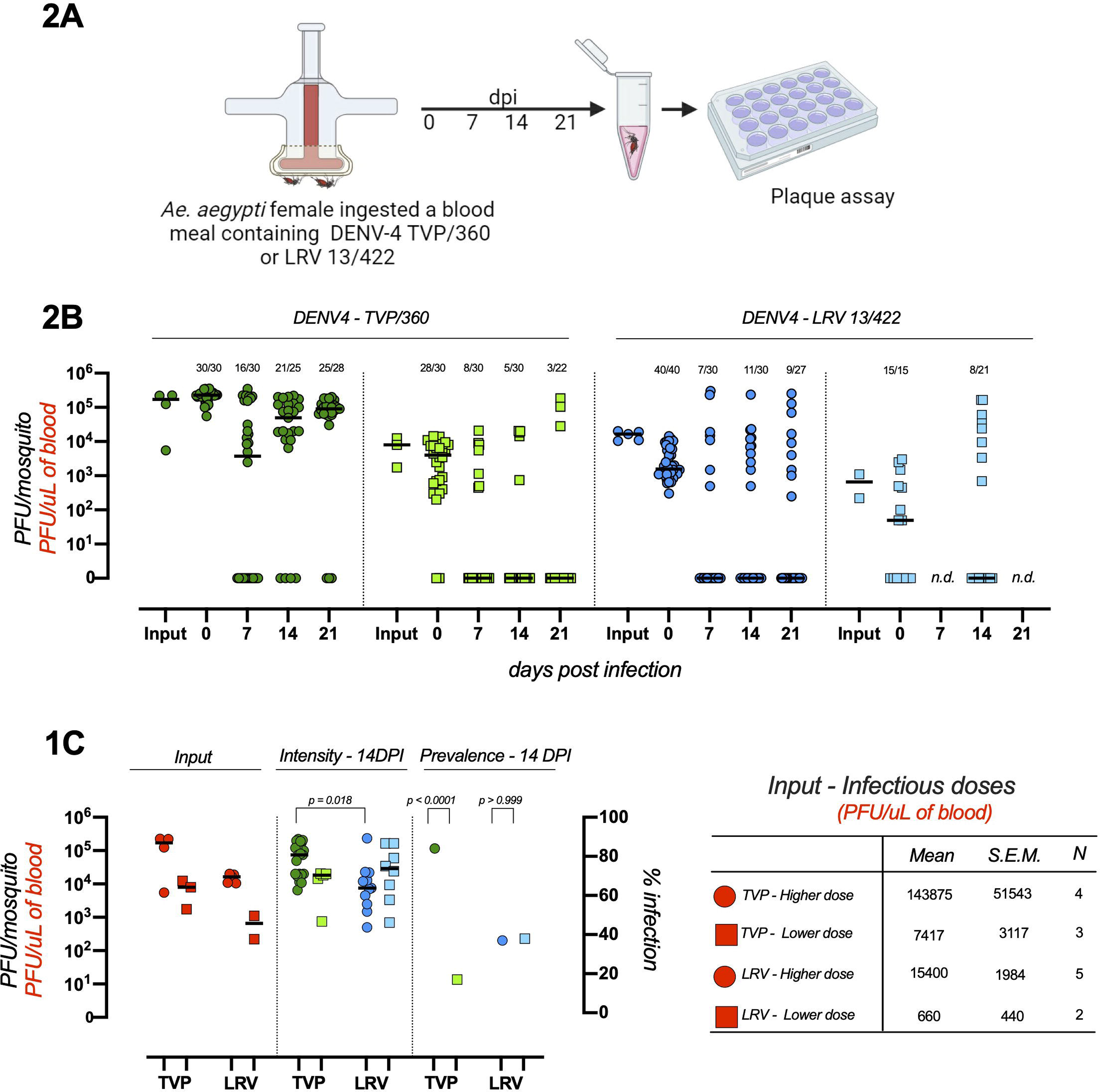
DENV4 TVP/360 and LRV 13/422 infection dynamics in *Aedes aegypti* mosquitoes. (A) Experimental design. Mosquitoes were fed with human blood containing different concentrations of two strains of DENV4 - TVP/360 and DENV4 - LRV 13/422. (B) Viral load was measured by plaque assay immediately following feeding (0), 7, 14, and 21 days post-infection (dpi). Each point represents 1 female mosquito. The number of samples tested is indicated above each group. Red dots represent the virus concentration in the blood offered to mosquitos in each replicate experiment. Bars in each column represent medians. This figure represents the sum of three to four independent experiments for each group. N.d.: - not determined. (C) Input is defined as the infectious particle concentration presented in the blood meal offered to mosquitoes (expressed as PFU/uL of blood - red text in the left Y axis). Intensity is defined as the number of infectious particles per mosquito measured at 14 days post-infection (PFU/mosquito in the left Y axis). Prevalence is defined as the percentage of infected mosquitos at 14 days post-infection (% infection in the right Y axis). Statistical analysis: Intensity - Kruskal Wallis test. Prevalence - Fischer’s exact test.

### The impact of different DENV4 strains on life-history traits of *Aedes aegypti*

We challenged mosquitoes with two doses of DENV4/TVP and DENV4/LRV and scored mosquito survival during the full life span of the population. Consistent with our previous results (Maraschin et al. 2023), the median time to death of *Aedes aegypti* did not differ between uninfected (mock) and infected mosquitoes, regardless of input viral doses and virus strain (Figure 3).

**Figure 3.**
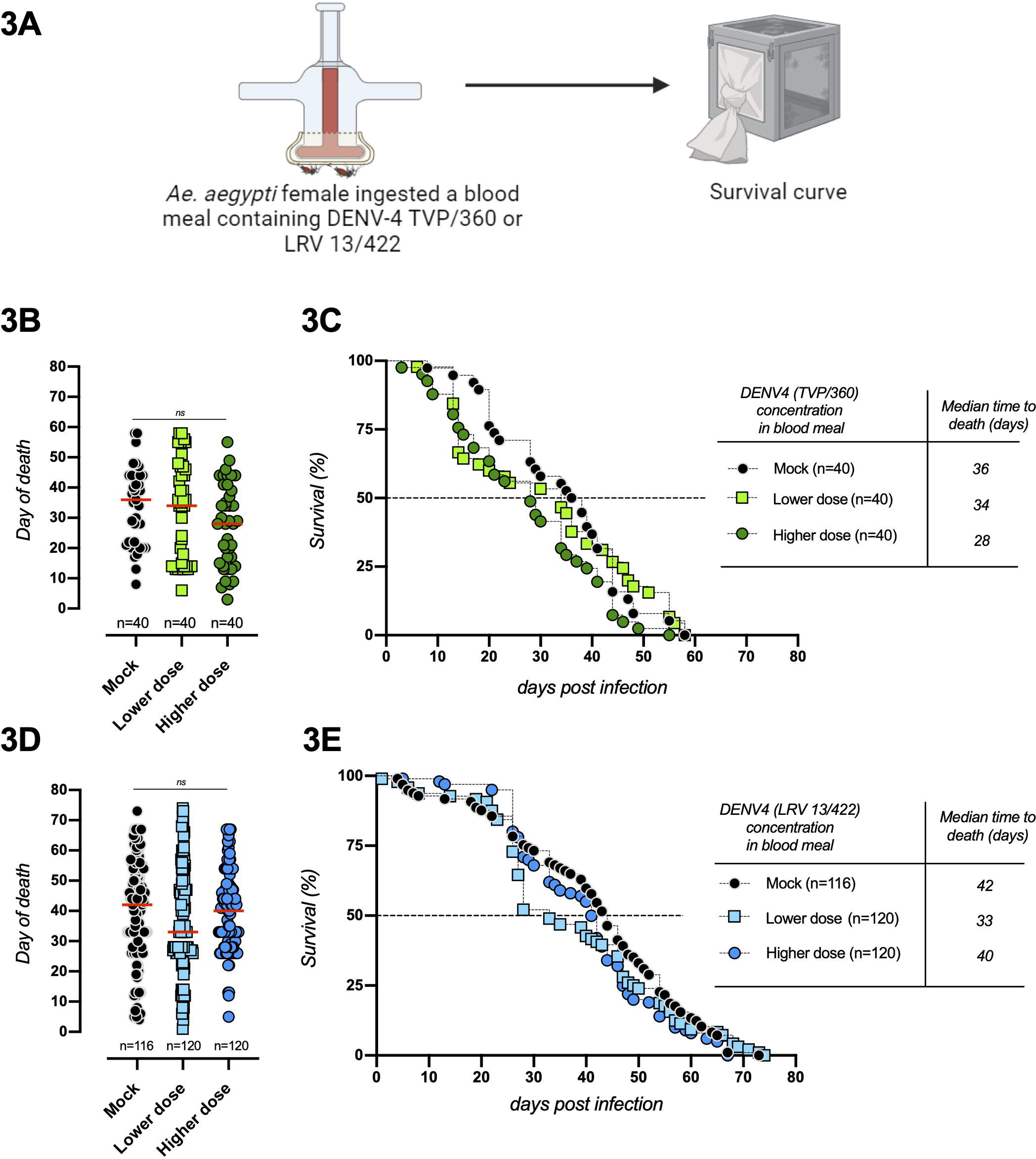
Survival curves of *Aedes aegypti* infected with DENV4 strain TVP/360 and DENV4 strain LRV 13/422. 3-4 days following adult emergence, females were fed with blood supplemented with 2 concentrations of DENV4 strains, as indicated in Figure 2C (Input - Infectious doses - PFU/uL of blood). Survival curves were performed at least twice in batches of 20 fully engorged females per cage. Survival was scored 6 times per week until all the mosquitoes died. Note that the survival of DENV4 (TVP/360) was already tested in much higher numbers in our previous work with similar results (Maraschin et al. 2023). Horizontal bars in 3B and 3D represent medians. Statistical analysis in Figure 3B and 3D - Kruskal Wallis test.

**Figure 4.**
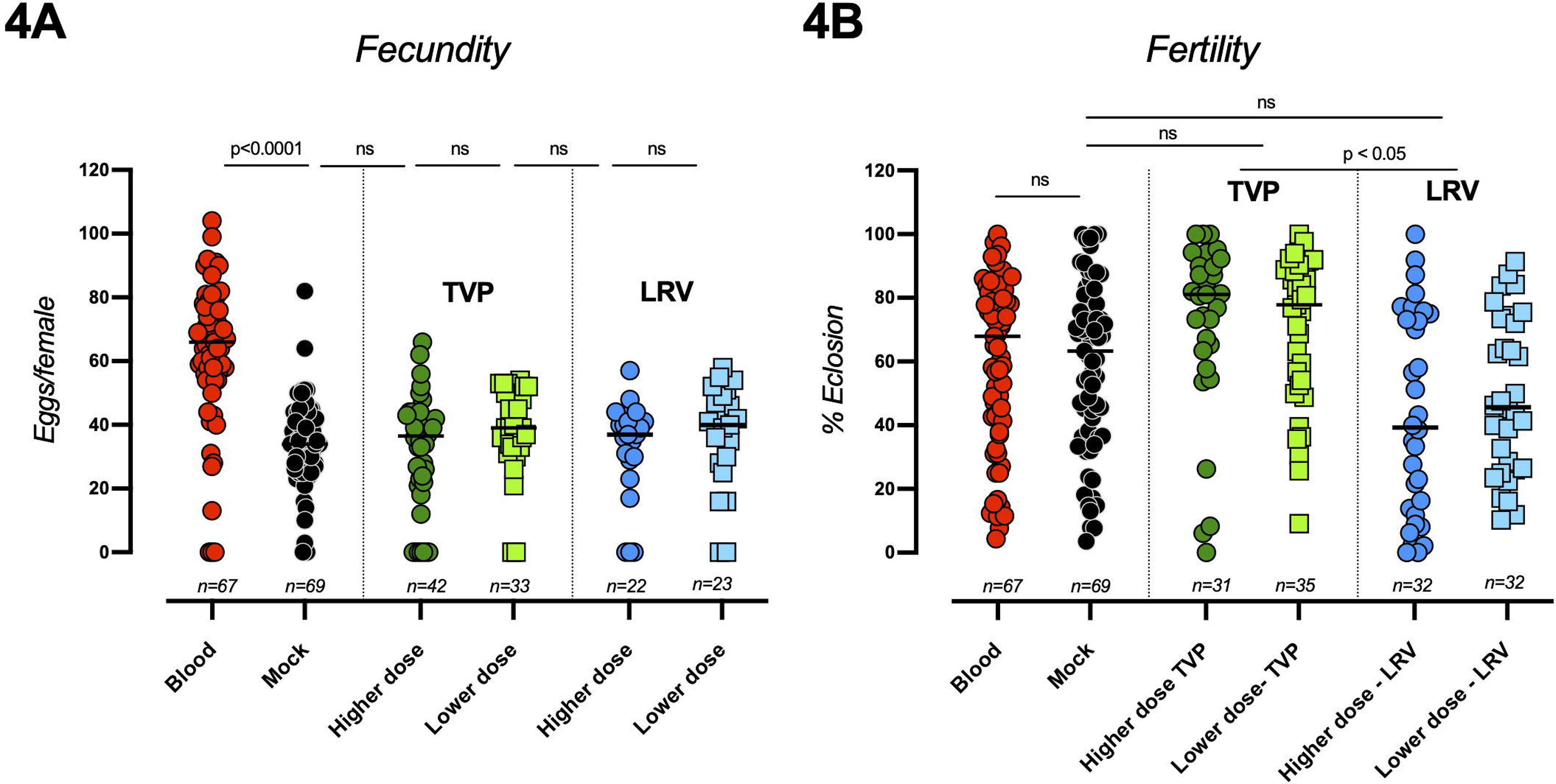
Fecundity and fertility of *Aedes aegypti* mosquitoes infected with DENV4 strains TVP/360 and LRV 13/422. Female mosquitoes were fed with human blood containing 2 concentrations of DENV4 strains, TVP or LRV, as indicated in Figure 2C (Input - Infectious doses - PFU/uL of blood). (A) Each point represents fecundity (eggs/female) measured through the individual oviposition of each mosquito. (B) Fertility represents the percentage of viable eggs laid by each mosquito individually measured 7 days post egg laying. Mosquitoes infected with DENV4 were compared with their respective controls: blood and mock (RBCs containing C6/36 cell supernatant instead of serum, as detailed in the materials and methods section). The number of samples tested is indicated at the bottom of each column. Experiments were conducted at least three times and present mean plus standard error. Statistical significance was determined using the Kruskal-Wallis test followed by Dunn’s multiple comparison test.

The impact of virus infection in parameters directly connected to vector fitness was evaluated following mosquito challenge with both DENV4 strains. Overall, feeding *Aedes aegypti* with blood supplemented with different concentrations of infectious particles did not alter the total number of eggs laid by each fully engorged female (Figure 4A). As detailed in materials and methods, we fed mosquitoes with human red blood cells (RBCs) supplemented with uninfected C6/36 cells supernatant (L-15 medium), referred to as “mock”, or RBCs supplemented with DENV4 infected C6/36 cells supernatant, referred as “higher dose” or “lower dose”. In Figure 4, “blood” represents whole human blood (RBCs + plasma), which has a higher total protein concentration than mock, as evidenced by a greater fecundity of *Aedes aegypti* (blood ∼70 eggs/female vs mock ∼35 eggs/female). Next, we scored fertility, defined as the number of eggs that hatched into L1 larvae 7 days following egg laying. Challenging mosquitoes with the clinical strain of DENV4 (LRV) resulted in a significant reduction in the percentage of eclosion (∼50%), irrespective of the dose tested (Figure 4B). This result suggests a lower adaptation of the LRV strain, which is consistent with its recent interaction with our colony mosquitoes (Red Eyes strain) instead of field mosquitoes circulating in southern Brazil (de Oliveira et al. 2023). On the other hand, we did not observe statistically significant differences in fertility with the laboratory-adapted TVP strain, consistent with the higher adaptability of this virus under laboratory conditions. The molecular basis of this phenotype is currently unknown. Interestingly, there is no difference in fertility between blood-fed and mock-fed mosquitoes, averaging around 70% eclosion, suggesting that females can optimize fecundity (egg production) according to the nutritional status of the meal to maximize fertility, similar to what has been described for desiccation stress in *Aedes aegypti* (Prasad et al. 2023).

*Aedes aegypti* dispersal involves its flight activity and is directly connected to vectorial capacity and arbovirus epidemics (Kraemer et al. 2015). We took advantage of a recently described method to assess the induced flight activity (INFLATE) (Gaviraghi and Oliveira 2020) and challenged *Aedes aegypti* females with the highest available doses of both strains of DENV4. The laboratory-adapted TVP/360 strain significantly enhanced the mosquito flight activity 24 hours after feeding when compared to “mock” and DENV4 - LRV 13/422 (Figure 5B), a time point where infection of the midgut epithelium is taking place. At 21 days post-feeding, an epidemiologically relevant time-point, where DENV4 has already infected the salivary glands and the mosquito is infectious (Salazar et al. 2007; Mayton et al. 2021), the INFLATE index was also significantly higher specifically in the laboratory-adapted TVP strain (Figure 5E). The clinical isolate did not induce alterations in the flight activity during the mosquito lifespan, compared to mock-infected *Aedes aegypti* (Figure 5B-E). Consistent with a reduction in mitochondrial respiration following a blood meal (Gonçalves et al. 2009), we observed an overall decline in INFLATE index 1 day post-feeding (Figure 5B) as reported by (Gaviraghi and Oliveira 2020), compared to all the other time-points. This reduction was independent of the DENV4 challenge. After the completion of blood digestion, at days 7, 14, and 21 days in our assays, mosquitoes were lean and lighter, which translated into higher INFLATE values compared to 1DPI (Figures 5C-E). At 21 days post-feeding, *Aedes aegypti* is already experiencing senescence, causing a reduction in the overall INFLATE index compared to 7 and 14 DPI (Figure 5E).

**Figure 5.**
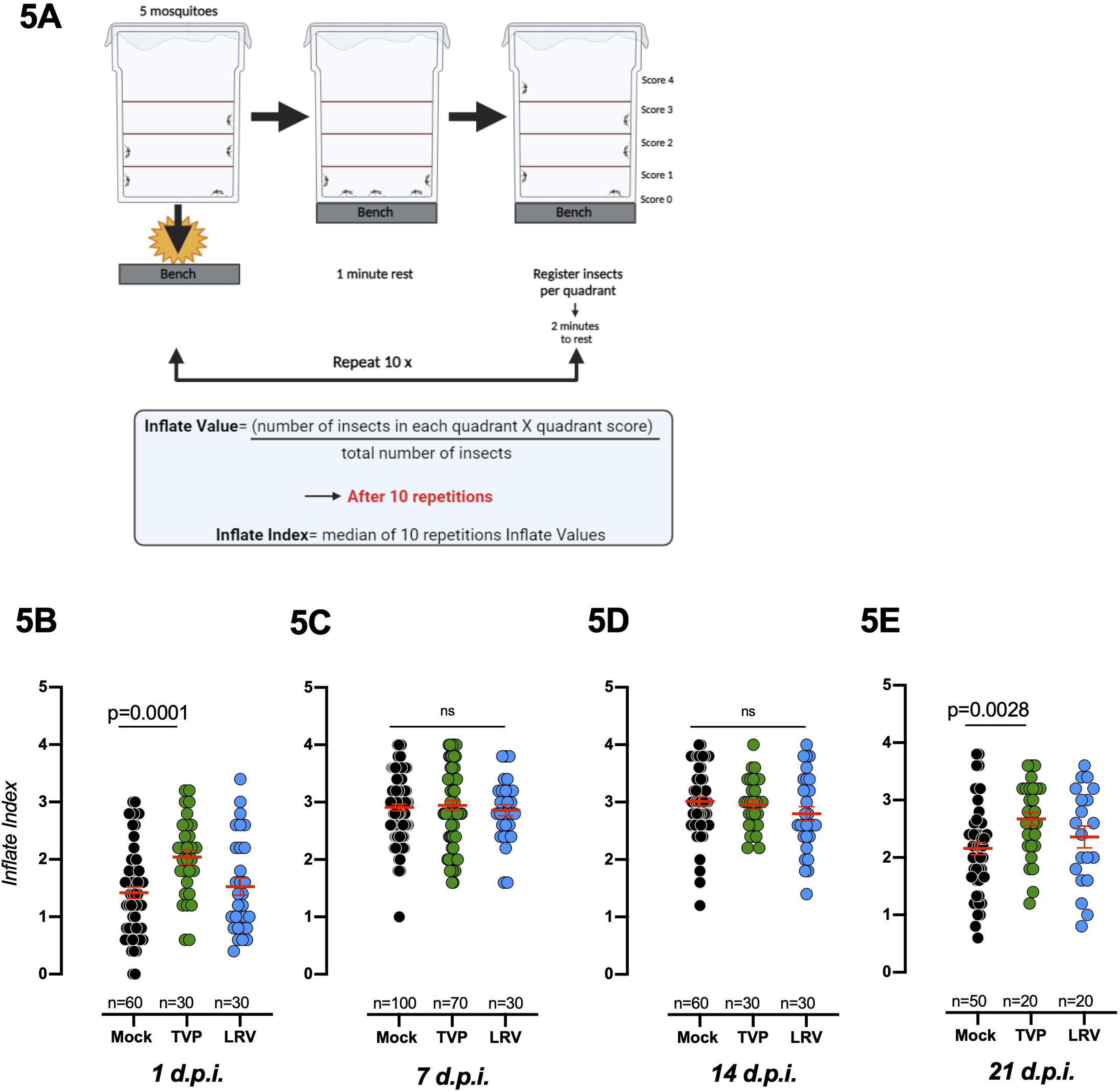
Comparative analysis of the induced flight activity of *Aedes aegypti* mosquitoes infected with DENV4 strains TVP/360 and LRV 13/422. (A) Experimental design of the INFLATE assay. See details in the methods section. Female mosquitoes were fed on blood containing the highest doses available for each strain - see Figure 2 (input). (B-C-D-E) The induced flight activity of each group was measured at 1, 7, 14, and 21 days post-infection (dpi). Mosquitoes challenged with DENV4 were compared with the mock/uninfected group. Each dot represents the inflate index (5 mosquitoes per cage tested 10 times in a row). The experiments were conducted at least 3 times, and statistical significance was determined by the Tukey multiple comparison test and ANOVA.

## 7 Discussion

Mosquitoes are tolerant to the microbes they transmit, such as dengue virus and plasmodium, a phenomenon that has been relatively well characterized, but poorly understood at the molecular level (Shaw et al. 2018; Oliveira 2020; Lambrechts and Saleh 2019; Shaw et al. 2022). In our previous work, we explored the impact of different arbovirus on mosquito mortality to establish a dose-response framework to study *Aedes aegypti’s* response to infections (Maraschin et al. 2023). A quantitative analysis of disease tolerance can be achieved by scoring how parameters relevant to host physiology vary during a gradient of infection (Louie et al. 2016; Torres et al. 2016; Gupta and Vale 2017). In this manuscript, we investigated the effect of two distinct DENV4 genotype II strains on mosquito life-history traits useful to describe organismal fitness and disease tolerance. We found that both DENV4 strains differentially impacted infection prevalence, female fertility, and induced flight capacity. Defining the molecular basis that underlines such phenotypes will be critical in efforts to mitigate mosquito-borne diseases based on vector control.

Dengue virus serotype 4 is one of the least studied dengue viruses. Although its presence in Brazil has been relatively limited (Nunes et al. 2012; Fang, Zhang, and An 2013; Ortiz-Baez et al. 2019; Bezerra et al. 2021), studies on the dynamics of different serotype circulation and emergence (Salles et al. 2018), as well as the occurrence of antibody-dependent enhancement of disease in humans (Dejnirattisai et al. 2010; Beltramello et al. 2010; Katzelnick et al. 2017), highlight the importance of examining the interaction between DENV4 and its main vector, the mosquito *Aedes aegypti.* This is particularly relevant since DENV4 has been shown to circulate undetected in mosquito populations without reported human cases (Boyles et al. 2020; Ayers et al. 2021), be vertically transmitted by mosquitoes (Torres-Avendaño et al. 2021) and displace DENV1, a common DENV serotype in Brazil (de Bruycker-Nogueira et al. 2018) when co-infecting *Aedes aegypti* (Vazeille et al. 2016). The strains used in our study exhibit 29 amino acid substitutions compared to DENV4 - 814669 (GenBank: AF326573.1), which was utilized as a reference (Figure 1 and Table 1). While the laboratory-adapted DENV4 - TVP/360 strain was closely related to the strain isolated in the Dominican Republic in 1981 (DENV4 - 814669), DENV4 - LRV 13/422, isolated in southern Brazil (Kuczera et al. 2016) exhibited the majority of amino acid substitutions; 24 in total (Tabel 1 and Supplementary Figure 1). Importantly, most of these mutations were found in critical structural and non-structural proteins that are essential for virus interaction and replication success within the mosquitoes, such as E, NS1, and NS5 (Table 1) (Erb et al. 2010; J. Liu et al. 2016; Y. Liu et al. 2017; Barrows et al. 2018; Saraiva et al. 2018; Wen et al. 2018; Besson et al. 2022; Cheng et al. 2021).

*Aedes aegypti* dispersal is a critical parameter to arbovirus infection in humans (Sedda et al. 2018; Leandro et al. 2024). We found that DENV4 - TVP/360, a laboratory-adapted strain, enhanced *Aedes aegypti* induced-flight capacity (INFLATE), a proxy for mosquito dispersal, specifically at an early (1DPI) and late (21DPI) time points (Figure 5B-E). This modulation was specific to the TVP strain since the recent clinical isolate (LRV) did not show differences in the INFLATE index compared to mock-fed mosquitoes (Figure 5B-E). It was previously shown that DENV infection enhances mosquito behaviours linked to flight (Lima-Camara et al. 2011; Tallon et al. 2020; Javed et al. 2024). Recently, immune activation by zymosan was shown to reduce *Aedes aegypti’s* induced flight capacity (Gaviraghi et al. 2024). The molecular and physiological basis of immune-induced flight phenotypes in *Aedes aegypti*, as well as its epidemiological implications, remain to be defined (Maire, Lambrechts, and Hol 2024).

The ability of dengue virus to infect and spread into the salivary glands of *Aedes aegypti* is highly dependent on virus and mosquito genotype interactions (Anderson and Rico-Hesse 2006; Fansiri et al. 2013; Fontaine et al. 2018; Dabo et al. 2024). Disease transmission is also influenced by environmental and ecological factors, such as temperature, nutrition, predation, humidity, and microbiota, among others (Gloria-Soria et al. 2017; Chandrasegaran et al. 2020). The virus infectious titer in a blood meal is a critical parameter in vector competence as it regulates mosquito infectivity and consequently, transmission potential, with higher doses leading to increased infection prevalence (% of infected mosquitoes) (Nguyet et al. 2013; Novelo et al. 2019; Johnson et al. 2024). The analysis of infection prevalence and intensity 14 days post-feeding (Figure 2C) reveals an interesting possibility regarding within-host strain adaptability in *Aedes aegypti*. DENV4 - TVP/360 is described as being laboratory-adapted and phylogenetically similar to the strain isolated in 1981 (Table 1 and Supplementary Figure 1). This strain was highly dependent on the input initial dose to establish infection (*p* < 0.0001 between TVP - higher dose vs TVP - lower dose - Prevalence at 14 DPI) (Figure 2C - TVP - green symbols). On the other hand, DENV4 - LRV 13/422, recently isolated in southern Brazil in 2013 and possessing several amino acid mutations in E, NS1, NS2A, and NS5 (Figure 1 and Table 1), did not exhibit an input dose-dependency to establish infection in our colony mosquitoes (*p* < 0.999 between LRV - higher dose vs LRV - lower dose) (Figure 2C - LRV - blue symbols - Prevalence at 14 DPI). This result might indicate that the recently isolated strain LRV 13/422 has evolved strategies to maximize infection prevalence over a wider range of input doses. In a real transmission setting, human viremia defines the virus input dose that mosquitoes will ingest during a blood meal (Long et al. 2019). In humans, plasma viremia increases following the infectious bite to a point where humans have enough circulating virus to transmit to uninfected mosquitoes (Duong et al. 2015), then viremia decreases rapidly, frequently over orders of magnitude every 24 hours (Vaughn et al. 2000; Carrington and Simmons 2014; Waggoner et al. 2016; Paz-Bailey et al. 2024). The prevalence data shown in Figure 2C suggests that the recent clinical isolate (LRV 13/422) has evolved strategies to infect mosquitoes that are less dependent on input doses/viremia, increasing the transmissibility window from humans to mosquitoes. This possibility reveals an interesting trade-off at the midgut infection barrier between TVP/360 and LRV 13/422, with higher input doses, that tend to occur in a shorter transmissibility window, favoring the TVP/360 strain, while the LRV 13/422 strain exhibits a lower infection prevalence over a wider period of infectiousness.

The contribution of different defense strategies, such as antiviral resistance or disease tolerance, during mosquito immune response to DENV is poorly defined ( Oliveira et al., 2020). While DENV load increases 100-1000 fold in mosquitoes during its spread from the midgut to the salivary glands (Salazar et al. 2007), *Aedes aegypti* does not experience significant fitness costs associated with chronic infection (Maraschin et al. 2023), although this may vary depending on different DENV serotypes, genotypes and mosquito populations (Keirsebelik et al. 2024). The RNAi machinery is considered the main antiviral pathway in *Aedes aegypti (Olmo et al. 2018)*, but the knockout or overexpression of its core gene, Dicer-2, revealed conflicting results regarding its actual contribution to mosquito immune resistance to arbovirus infection and vector competence (Dong et al. 2022; Merkling et al. 2023; Samuel et al. 2023). In Figure 2C we show that despite a marked difference in prevalence at 14DPI between the LRV 13/422 and TVP/360 strain depending on the initial DENV4 input dose, infection intensity (Figure 2C - middle panel) was not influenced. This result suggests that immune resistance plays a minor role in the DENV4 replication cycle within the mosquito and unknown factors controlling the midgut infection barrier might be more important to mosquito infection and, therefore, disease transmission.

In summary, we described that two different strains of DENV4 genotype II differentially impacted *Aedes aegypti* infection dynamics, fertility and flight capacity. The determination of how arbovirus infection affects life-history traits of insect vectors is critical for the development of strategies to fight mosquito-borne diseases (Achee et al. 2019).

## 9 Authorship contribution

Mariana Maraschin: data collection, data analysis, writing.

Diego Novak: data collection, data analysis.

Valdorion José Klein Junior: data collection, data analysis.

Lucilene W Granella: data collection.

Luiza J Hubner: data collection.

Athina R Madeiros: data collection.

Tiago Graf: data analysis, writing.

Guilherme Toledo-Silva: data analysis.

Daniel S Mansur: data analysis, funding acquisition, writing, supervision.

Jose Henrique M. Oliveira: conceptualization, data analysis, funding acquisition, writing original draft, supervision.

