## Supplementary figures and images for "A Laboratory-Adapted and a Clinical Isolate of Dengue Virus Serotype 4 Differently Impact Aedes aegypti Life-History Traits Relevant to Vectorial Capacity"

### Supplementary Figure 1 - FINAL.jpg

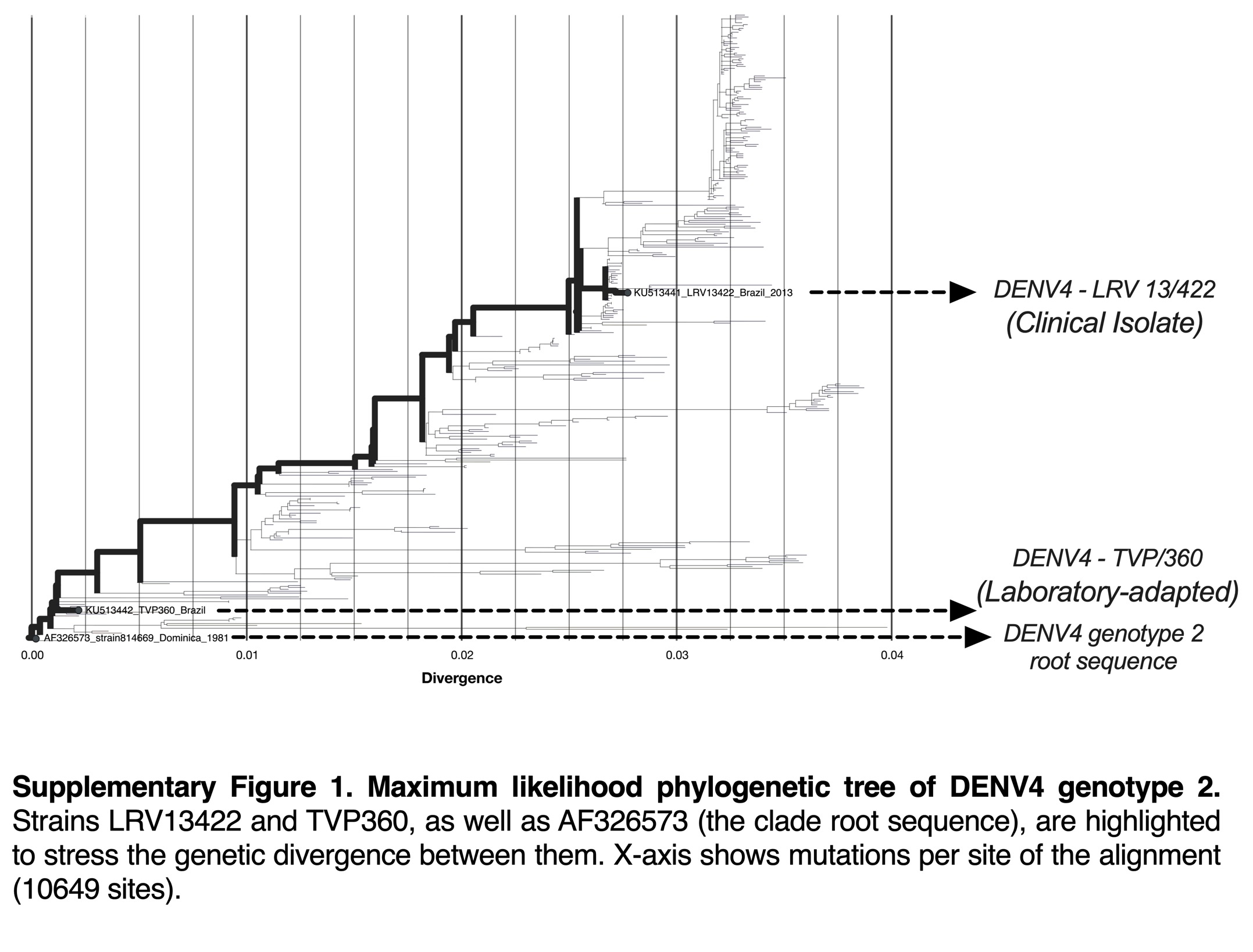
